# Development of an optogenetic gene expression system in *Lactococcus lactis* using a split photoactivatable T7 RNA polymerase

**DOI:** 10.1101/2024.01.05.574370

**Authors:** Aleixandre Rodrigo-Navarro, Manuel Salmeron-Sanchez

## Abstract

Cellular processes can be modulated by physical means, such as light, which offers advantages over chemically inducible systems with respect to spatiotemporal control. Here we introduce an optogenetic gene expression system for *Lactococcus lactis* that utilizes a split T7 RNA polymerase linked to two variants of the Vivid regulators. Depending on the chosen photoreceptor variant, either ‘Magnets’ or ‘enhanced Magnets’, this system can achieve either high protein expression levels or low basal activity in the absence of light, exhibiting a fold induction close to 30, rapid expression kinetics, and heightened light sensitivity. This system functions effectively in liquid cultures and within cells embedded in hydrogel matrices, highlighting its potential in the development of novel engineered living materials capable of responding to physical stimuli such as light. The optogenetic component of this system is highly customizable, allowing for the adjustment of expression patterns through modifications to the promoters and/or engineered T7 RNA polymerase variants. We anticipate that this system can be broadly adapted to other Gram-positive hosts with minimal modifications required.

## Introduction

*Lactococcus lactis* is an important gram-positive probiotic bacterium that has been the subject of intense research during the last few decades and has a generally regarded as safe (GRAS) status. Besides its widespread use in the food industry as a starter culture for the manufacturing of dairy products like buttermilk and cheese, it has gained traction as a live vector for mucosal vaccine delivery(*1*) and recombinant protein production without the disadvantages of *E. coli* such as the contamination of the recombinant proteins with lipopolysaccharides that can elicit immune response and inflammation, and the presence of robust protein secretion pathways in *L. lactis* compared to *E. coli*(*2*). *L. lactis* has been used recently in tissue engineering applications and in engineered living materials (ELMs) (*3–5*), a nascent field where a new class of materials based on an inert matrix and a living component, usually engineered using synthetic biology, combine together to produce “smart” functional materials with capabilities that surpass the current state-of-the-art. ELMs are able to respond to environmental physicochemical cues, namely electromagnetic and/or mechanical signals, chemical species and gradients, pH and others. The ELM must be able to integrate and respond to those signals and dynamically switch between different states, responding with output signals ranging from the production of proteins/chemicals to changes of in the properties of the ELM itself, phenotypical changes of the living component and so on. The implications and possible use cases for ELMs are broad and include biosensors, microbial cell factories and bioreactors for the production of high added value products, tissue engineering or agriculture construction or bioremediation.(*6*, *7*)

The use of bacteria as the living component in ELMs for biomedical applications requires a certain degree of control over how genes, and thus proteins or small molecules, are modulated to produce therapeutic effects in the form of the expression of chemicals and/or proteins. *Lactococcus lactis* is an organism with a limited genetic toolbox in comparison to other model species such as *Bacillus subtilis* or *Escherichia coli*, so the availability of regulated gene expression systems was limited to chemically/environmentally inducible cues. Several examples of inducible systems are described in the literature, which use environmental stimuli such as the transition to low pH(*8*), stress conditions(*9*), zinc ions(*10*, *11*), xylose(*12*), agmatine(*13*), phages(*14*, *15*) or nisin through the nisA promoter(*16*), this system being the most popular due to the high expression levels achievable while maintaining a tight control over basal gene expression. However, even if chemically or environmentally inducible gene expression systems, especially the food-grade ones, are useful for biotechnological applications, its utility is limited in terms of spatiotemporal control, due to the difficulty to induce only a part of the system. Moreover, gene expression deactivation requires the removal of the inducer from the system, which can be costly and cumbersome. This behaviour poses a disadvantage when working with rheostats, where the gene expression cannot be switched off in a straightforward manner.

Here we show the development of a novel optogenetic system for *L. lactis*. The basis for our system is the single unit DNA-dependent RNA polymerase from the *E. coli* bacteriophage T7 (T7RNAP)(*17*, *18*), fused to an engineered variant(*19*) of the fungal *Neurospora crassa* homodimerizing photoreceptors Vivid (VVD)(*20*), using an approach similar to a previously described *E. coli* system(*21*), and another variant using the same rationally designed and improved photoreceptors(*22*). T7RNAP is a highly processive single-subunit polymerase with a high selectivity for its cognate T7 promoter and the ability to produce very long transcripts(*23*). It founds widespread use as a molecular biology tool for heterologous gene expression. T7RNAP-based expression systems have been successfully translated to mammalian cells (*24*), yeast strains like *Saccharomyces cerevisiae*(*25*) and *Pichia pastoris* (*26*), plants like tobacco (*27*), and other bacterial species, either gram-negatives like *Vibrio natriegens*(*28*, *29*) or gram-positives such as *Bacillus subtilis*(*30*) and *Lactococcus lactis*(*31*).

## Results

The basis for the present optogenetic gene expression system is the phage T7 RNA polymerase harbouring the R632S mutation, which has been shown to reduce its processivity and cytotoxicity(*32*). It has been also shown that the T7RNAP can be split at different positions and reconstituted keeping its activity(*33–36*).

The present optogenetic system is based on the phage T7 RNA polymerase split at the residue 563, with each fragment fused to an engineered variant of the homodimerizing Vivid (VVD) photoreceptors from the filamentous fungi *Neurospora crassa*, called Magnets(*19*), or to an improved version called “enhanced Magnets” or eMags(*37*), in both cases through a flexible glycine/serine GGSGG linker. The T7RNAP can be split in different positions and reconstituted by dimerization while keeping its activity. Inspired on a previous work(*21*), our goal was to split the T7RNAP and fuse the C- and N-terminal ends to either the nMagHigh1 and pMagFast2 variants of Magnets photoreceptors or its “enhanced” (eMagA and eMagB, the equivalents of the nMagHigh1 and pMagFast2) variants using a flexible linker, in a way that exposure to 450 nm light heterodimerizes the photoreceptors, bringing together the two polymerase subunits and restoring its activity.

We split the T7RNAP at the position 563 and fused the N-terminal end to either nMagHigh1 or eMagB Magnet variant and the C-terminal end to the pMagFast2 or eMagA variant. The R632S mutation in the T7RNAP(*32*) reduces its processivity and thus its cytotoxicity, while keeping its activity, even if previous studies have shown that mutations to this region of the polymerase can have an effect on its processivity(*38–40*).

Since the copy number of active T7RNAP and consequently the derived protein overexpression can have a profound effect on the bacterial metabolic machinery and ultimately a cytotoxic effect(*18*, *41*, *42*), we screened constitutive lactococcal promoters in an attempt to express the polymerase at low levels. We used a library of synthetic promoters with variable potency(*43*) and constructed a series of plasmids where superfolder GFP (sfGFP) (*44*), a rapidly folding and robust version of GFP that works especially well in *L. lactis*, was expressed under the control of few of them (figure 1A). We wanted to estimate the potency of each promoter in terms of sfGFP fluorescence per CFU, assuming that the only factor affecting the expression levels was promoter, since we used the same RBS sequence in each experiment. The fluorescence values of individual cells with sfGFP expressed under the control of variable potency promoters were measured using flow cytometry (figures 1C and 1D). The log-log graph of promoter intensity, expressed as β-galactosidase activity in the original work, versus sfGFP fluorescence measured as the median of the fluorescence intensity histograms obtained in flow cytometry, gave an approximate linear correlation. We used promoters with relative potencies of 0.3, 5.1, 50 and 1000 (CP03, CP5, CP50 and CP1000 from now on) in terms of β-galactosidase activity units.

**Figure 1.**
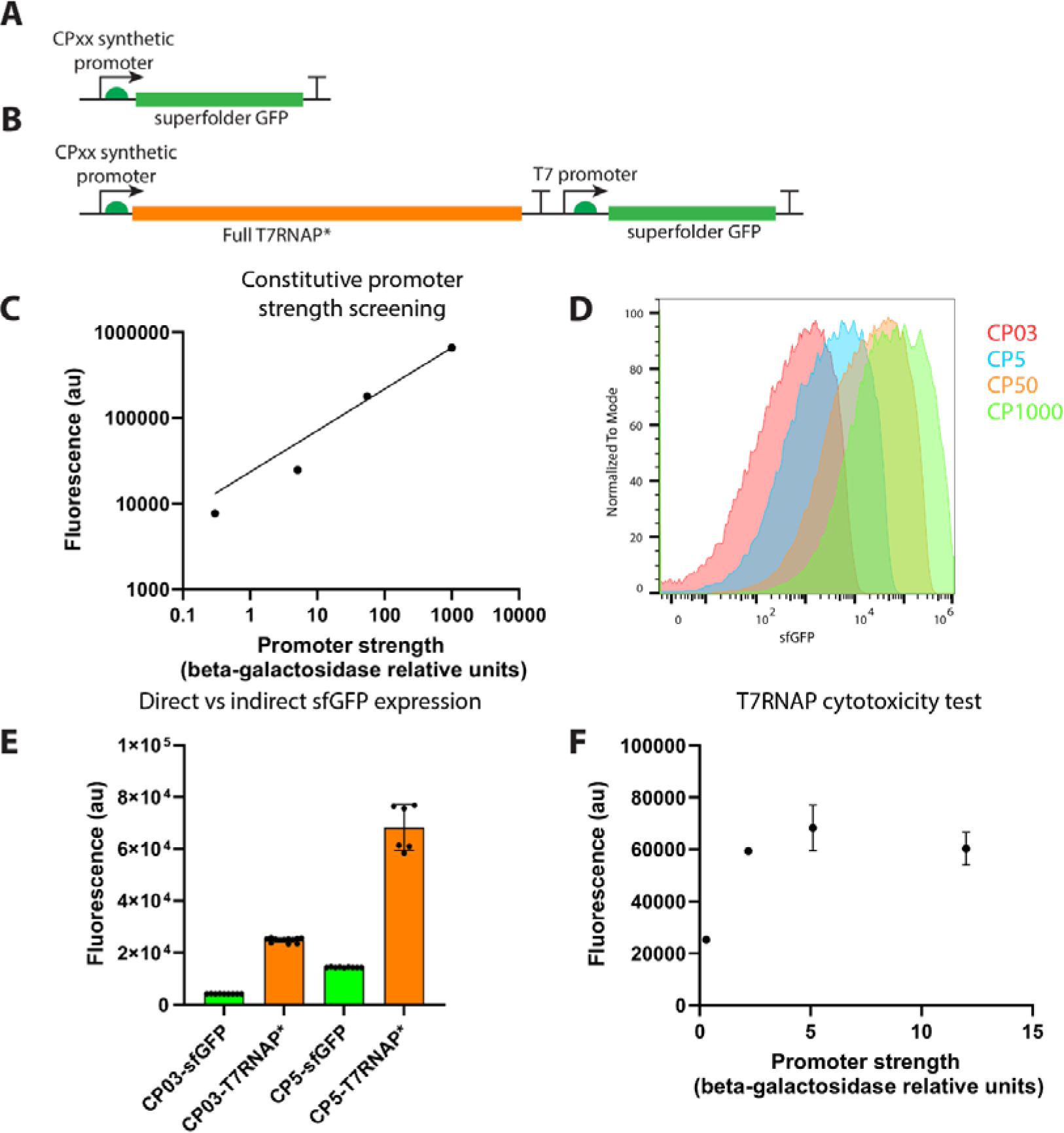
A) Schematic for the plasmids used to screen the potency of the CPxx synthetic promoters, where the sequences of the different promoters were inserted upstream the T7g10 5’-UTR and RBS to drive the expression of sfGFP, that was later quantified using either fluorometry or flow cytometry. B) Schematic of the plasmids used to screen the different promoters for the expression of sfGFP under the control of the full T7RNAP. Four synthetic promoters were used, CP03, CP2, CP5 and CP12, and sfGFP was expressed under the control of the T7 promoter and the *L. lactis*-optimized 5’-UTR and RBS. C) Log-log graph of the fluorescence intensity, measured with a fluorometer, of *L. lactis* strains transformed with plasmids expressing constitutively sfGFP under the control of the CP03, CP5, CP50 and CP1000 promoters. The theoretical strength of the promoter is closely followed by the data, showing a monotonic increase in fluorescence as the promoter strength was higher. D) Flow cytometry histograms of the same cells used in D, showing in this case the values for the sfGFP fluorescence. There is a unimodal distribution of the intensities, following closely the results from graph D. E) Fluorescence intensity measured in a fluorometer of *L. lactis* strains transformed with plasmids either expressing sfGFP under the direct control of the CP03 and CP5 promoters or using the same promoters to express the full T7RNAP that would later drive the expression of sfGFP under the control of the T7 promoter. As shown in the graph, the T7 polymerase has an amplificatory effect with an average fold change of 5.33. F) In order to determine if there was a toxic effect of the T7RNAP dose on the cell viability and thus sfGFP expression levels, two more promoters were cloned upstream the T7RNAP that would drive sfGFP expression under the T7 promoter, namely CP2 and CP12. As it can be seen in the graph, after CP2 the signal reaches a plateau. We observed that the bacteria would grow normally without diminishing the final biomass values (data not shown), suggesting that even higher strength promoters would not be toxic for the cells. We thus decided to use either the CP2, CP5 or CP12 promoters to drive T7RNAP constitutive expression to construct the final optogenetic plasmids.

The next step was to confirm the suitability of the T7RNAP as an orthogonal expression system in *L. lactis* NZ9000, as previously described(*31*). The T7RNAP gene, including the R632S mutation, was amplified using PCR from the pN565 plasmid(*45*) and subcloned under the control of four different synthetic promoters with varying potencies (CP03=0.3, CP2=2.2, CP5=5.1 and CP12=12 relative β-galactosidase units), to assess how the expression at different doses would affect in the expression of genes under the control of the T7 promoter (figure 1B).

As shown in figure 1E, the use of the same promoter to express sfGFP directly, or indirectly via T7RNAP, yielded different values, as expected from the amplificatory effect of the T7RNAP. We observed an approximate fold change value of around 5.9, with a slight dependency on the promoter. We also wanted to test if the expression of the T7RNAP at higher levels would be toxic to the cells, but we observed (figure 1F) that promoters with a strength equal or higher than 2.2 (CP2, CP5 and CP12) resulted in similar sfGFP fluorescence values, reaching an apparent plateau. It might be possible that the lower processivity of the T7RNAP variant with the R632S mutation makes it less toxic than the full native T7RNAP, although we did not test this hypothesis.

It is worth noting that during the initial attempts to clone the T7RNAP under the control of the synthetic promoters, we observed a G213V mutation that further reduced the activity of the polymerase (Supplementary figure 1A). This mutation was later reversed, and the polymerase returned to its baseline activity, with a fold change of approximately 5.9 between indirect sfGFP expression through T7RNAP compared to direct expression (Supplementary figure 1B). Even if not strictly relevant for this work, this result might be useful for T7RNAP-driven systems that require even lower expression values while still benefiting from the orthogonality of this polymerase relative to the native translation machinery of *L. lactis*.

Once T7RNAP’s activity and lack of toxicity was confirmed, it was split at the residue 563(*21*), fusing the nMagHigh1 variant of the engineered Vivid photoreceptor to the N-terminal fragment of the T7RNAP, and the C-terminal end to the pMagFast2 variant using a GGSGG flexible linker (figure 2A). To confirm that both the N- and the C-terminal fragments T7RNAP fused to nMagHigh1 and pMagFast2 respectively lacked transcriptional activity, we constructed two plasmids where either fragment was expressed under the control of the CP12 synthetic promoter. The sfGFP gene was cloned under the control of the T7 promoter.

**Figure 2.**
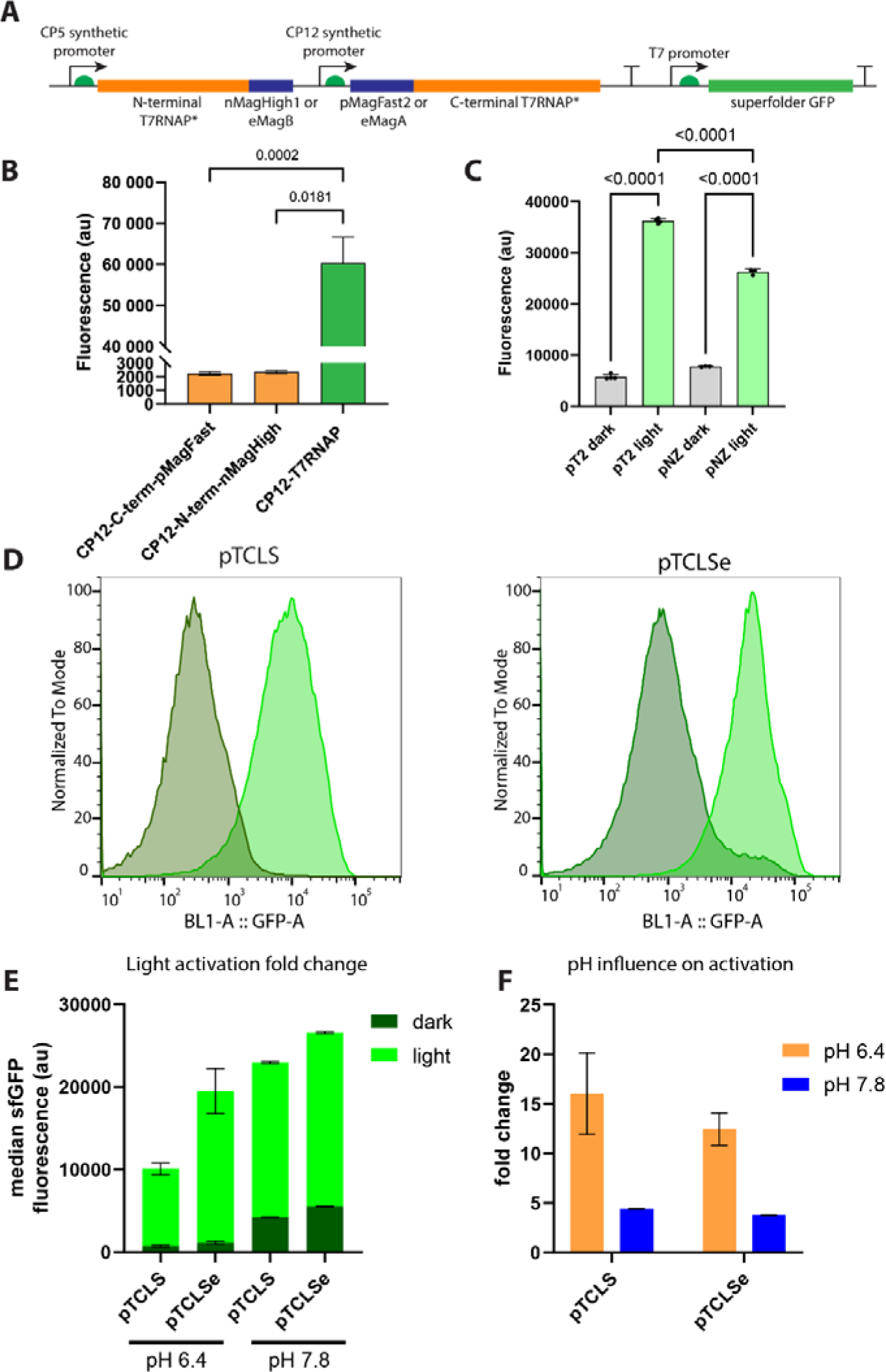
A) Schematic diagram of the construction of the pCLS, pTCLS and pTCLSe plasmids. Two synthetic promoters, CP5 and CP12, control the constitutive low-level expression of the two fragments of the T7RNAP fused to either nMagHigh1/pMagFast2 or eMagA/eMagB. The superfolder GFP (sfGFP) gene is located downstream the T7 promoter and the 5’-UTR and RBS optimized for *L. lactis*. B) To determine if the split T7RNAP has any residual activity, either the N-terminal or C-terminal fragments fused to eMags or nMagHigh1/pMagFast2 were knocked out from the plasmid, transformed in *L. lactis* and the sfGFP fluorescence was measured in a plate reader and compared to the fluorescence of the sfGFP expressed from the full T7RNAP. The CP12 synthetic promoter was used to control the T7RNAP expression levels. The fluorescence observed with the full T7RNAP was significantly higher than each fragment by itself, that was comparable with a negative control consisting of wild type *L. lactis* (data not shown). C) The first attempt to construct the optogenetic T7RNAP expression system was performed in a modified pNZ8123 plasmid with a pSH71 origin of replication (rolling circle replication) but recombination events forced the cloning of the optogenetic system in the pT2NX plasmid, with a pAMβ1 origin of replication and a theta (θ) replication mechanism. This change led to an increased structural stability and an improved fold change between the inactive and active states. D) *L. lactis* transformed with pTCLS or pTCLSe were exposed to a blue LED (450 nm) with an intensity of 300 μW/cm^2^ for 5 hours, and the GFP fluorescence was measured with an Attune NxT flow cytometer. E) The pTCLSe plasmid shows increased gene expression levels, translated here as GFP fluorescence, but at the same time shows higher basal expression levels. F) The pTCLS plasmid shows here a slightly improved fold change between the dark and activated states but lower maximum expression levels.

As expected (figure 2B), T7RNAP fragments fused to the photoreceptors showed negligible sfGFP fluorescence compared to the plasmid with the full-length T7RNAP, suggesting an absence of polymerase activity. For the optogenetic expression system we chose a combination of the nMagHigh1 and pMagFast2 photoreceptor variants(*19*) due to their improved dissociation kinetics and acceptable heterodimerization efficiency. Specifically, the nMagHigh1 and pMagHigh1 variants display faster dissociation kinetics (t_1/2_ = 4.2 min vs. 1.8 h for the original pMag/nMag pair) but with an approximately 51% lower dimerization efficiency. However, the combination of nMagHigh1 and pMagFast2 variants increases dimerization efficiency by 760% compared to pMagFast2 and nMagFast2 while retaining fast dissociation kinetics.

We firstly constructed a single plasmid optogenetic system named pCLS, with the split T7RNAP fused to the Magnets and the T7 promoter driving the expression of sfGFP, using the pNZ8123 backbone. This plasmid is a gram-positive broad-range expression vector originally used for the nisin expression system and features a pSH71 origin of replication with a rolling-circle replication mechanism and chloramphenicol acetyltransferase antibiotic resistance gene for selection. The original P_NisA_ promoter and other elements were removed from the plasmid and the split N-terminal-nMagHigh1 and C-terminal-pMagFast2 fragments of the T7RNAP, under the control of the CP5 and CP12 promoters respectively, a modified T7 promoter optimized for *L. lactis*(*31*) followed by the sfGFP gene and the T7 terminator were subcloned using a mixture of synthetic gene fragments and PCR products via Gibson assembly. The first experiments showed light-activated sfGFP expression, demonstrated as a clear difference in the sfGFP fluorescence in bacterial cultures grown overnight either in vessels protected from light, or in ambient light with an estimated 450 nm irradiance of around 70-75 μW/cm^2^ (figure 2C), demonstrating the viability and efficiency of this system. After few experiments using consecutive subcultures of *L. lactis*-pCLS we noticed that the bacteria lost its ability to express sfGFP under the control of the optogenetic T7RNAP. The pCLS plasmid was found to be noticeably smaller, and after sequencing it, we found deletions and recombinations near to or in the pMagFast2/nMagHigh1 genes. Both genes share a homology of 98.2% since they are the same protein with few mutated residues. It was immediately obvious that the rolling circle replication mechanism of pCLS was the most probable cause of these recombinations. Further experiments showed that after three to five subcultures, *L. lactis*-pCLS lost in all cases its ability to express sfGFP under the influence of blue light (data not shown).

To overcome this issue, we decided to codon-optimize pMagFast2 and nMagHigh1 to reduce the homology, and subcloned the optogenetic system in the pT2NX plasmid, a derivative of the plasmid pT1NX(*46*) with the erythromycin resistance methylase (*erm*) gene replaced by a chloramphenicol acetyltransferase gene (own work). The pT2NX plasmid has the origin of replication of the pAMβ1 plasmid, with a theta replication mechanism, allowing the cloning of larger inserts and an improved structural stability(*47*). Light activation experiments showed improved sfGFP expression under ambient light and a lower basal activity compared to pCLS (figure 2C, where pNZ refers to the pNZ8123 plasmid with a pSH71origin of replication and pT2 to the pT2NX plasmid with a pAMβ1origin of replication). The new plasmid was designated as pTCLS and used for subsequent experiments. Before continuing with the experiments, we checked the sfGFP photobleaching sensitivity under our experimental conditions, that is, λ=470 nm light with an irradiance up to 300 μW/cm², and how that would affect the results. Superfolder GFP undergoes photolysis when exposed at blue light (445 nm) at intensities over 306 mW(*48*), which is 3 orders of magnitude above our experimental conditions. We observed (Supplementary figure 2) that sfGFP fluorescence did not show a quantifiable difference even after irradiation for up to 5h at 300 μW/cm^2^, making in our criteria sfGFP a valid reporter protein to evaluate the system.

*L. lactis*, as a homofermentative anaerobic lactic acid-producing strain, metabolizes glucose to lactic acid using a glycolytic homofermentative pathway in anaerobic conditions, being glycolysis the main source for the biosynthesis of ATP in these conditions. As a result, lactic acid lowers the environmental pH to values around 5 to 5.5, rendering the T7RNAP inactive(*49*). A simple and straightforward way to minimize this problem was the supplementation of L-arginine in the medium as an alternative carbon source taking advantage of the lactococcal arginine deiminase (ADI) pathway, that metabolizes arginine to L-ornithine, NH_3_ and CO_2_ as a complementary ATP source(*50*), raising the pH to approximately 6.5-7 due to the effect of the ammonium cation that counteracts the acidity of the lactic acid. As a result of the increased pH value, the T7RNAP recovers its activity. A combination of 0.5% glucose and 0.5% arginine yielded a final pH of around 6.4, where the T7RNAP is active again, and L-arginine alone, at concentrations ranging from 0.5 to 1% yielded pH values of around 7.8 to 8 respectively, while keeping an adequate biomass production (supplementary figure 3). In this way, we made sure that the T7RNAP is active, and it is possible to assess the influence of light irradiation on the magnets to evaluate the system.

During the construction of the system based on the original Magnets, it came to our attention the development of an enhanced variant(*22*) that aimed to tackle the limitations of the nMag/pMag variants, namely the need of a 24 h preincubation step at 28°C to allow for the folding and maturation of the receptors, the necessity of concatemerization to improve their dimerizing efficiency, and their inability to dimerize at 37°C. We replaced the nMagHigh1 with eMagB and pMagFast2 with eMagA using Gibson assembly, obtaining the plasmid pTCLSe as a result.

In both pTCLS (nMagHigh1/pMagFast2) and pTCLSe (eMagB/eMagA, figure 2A) the N- and C-terminal fragments of the T7 RNA polymerase were expressed under the control of the CP5 for the N-terminal and CP12 promoters for the C-terminal fragments respectively, keeping the flexible glycine/serine linker, while sfGFP expression was under the control of the T7 promoter adapted to *L. lactis*.

To assess the influence of the original and enhanced photoreceptors on the sfGFP expression under blue light, *L. lactis*-pTCLS and *L. lactis*-pTCLSe cultured in M17 medium supplemented with 0.5% glucose and 0.5% L-arginine (GAM17C) or 0.5% L-arginine only (AM17C) and 10 μg/mL chloramphenicol in vessels protected from light (“dark” condition in the figures) or irradiated with blue LEDs (“light” condition in the figures) at 300 μW/cm² for 5h.

Immediately after the experiment, the bacterial cultures were cooled down to 4°C, centrifuged and fixed. Since *L. lactis* is a species that grows in chains, the fixed cultures were homogenised in a Qiagen TissueLyser apparatus at 50 Hz for 2 minutes to disrupt the chains. The cultures were analysed in an Attune NxT flow cytometer. Cells cultured in absence of light (figures 2C and 2D) showed lower fluorescence values compared to cells cultured with blue light irradiation. We observed that the pTCLS plasmid showed an increased fold change between the basal and activated states (figures 2E and 2F) compared to the pTCLSe plasmid when using GAM17C. This fold change was lower in cells cultured in AM17C, but the overall protein expression levels were significantly higher, with a higher final value of the medium and consequently a more active T7RNAP, and again the pTCLSe plasmid yielded higher expression levels compared to pTCLS.

We are aware that depending on the application, it might be desirable to achieve an overall higher protein expression level, for which the pTCLSe plasmid and/or supplementing the culture medium with L-arginine would be more suitable, while in other scenarios a higher dynamic expression range might be more convenient, for which the pTCLS plasmid combined with pH adjustment via using variable concentrations of L-arginine could be a better strategy. We did not test intermediate conditions with variable concentrations of glucose and L-arginine, but it seems logical that controlling the pH of the medium could be a sensible approach.

Next, we examined the effects of irradiance intensity on photoreceptor heterodimerization and resulting gene expression levels. According to literature, heterodimerization of Magnets reaches a maximum at around ≈300 μW/cm² in *E. coli* (*21*), so we tested up to this light intensity. We cultured *L. lactis*-pTCLS and *L. lactis*-pTCLSe strains overnight in opaque vessels protected from light using static conditions, in either GAM17C or AM17C medium. The following day, we diluted the cultures 1/5 into fresh GAM17C or AM17C medium prewarmed to 30°C and then exposed them to blue light at intensities of 0, 10, 40, 80, 100, 200 and 300 μW/cm² for a total of 5 hours. Irradiance intensity control was accomplished using an in-house made device which consists of an Adafruit Feather M4 development board with an ATSAMD51 chipset connected to a TLC5947 PWM 24-channel LED controller programmed in Python and calibrated to output blue light from 0 to 2000 μW/cm², which is the maximum irradiance offered by the blue LEDs.

In *L. lactis*-pTCLSe bacteria cultured at pH ≈6.4 (medium supplemented with glucose/arginine) sfGFP expression increased monotonically to up to 300 μW/cm² without plateauing, while in the medium supplemented with only arginine (pH ≈7.8) sfGFP expression reached a higher plateau starting at the lowest irradiance of 10 μW/cm². In the case of the pTCLS plasmid, the expression levels at lower pH ≈6.4 reached a maximum after 80 μW/cm² with overall lower expression levels and showed a similar behaviour at higher pH compared to the pTCLSe plasmid, reaching a higher plateau from 10 μW/cm². It is interesting to note that the pTCLSe plasmid yielded similar expression levels at pH ≈6.4 at the maximum tested irradiance, even with decreased T7RNAP activity, than the pTCLS plasmid at pH ≈7.8 (figure 3C). This is a strong indicative that the eMags dimerize with higher efficiency under the same light intensity, as expected from the literature.

**Figure 3.**
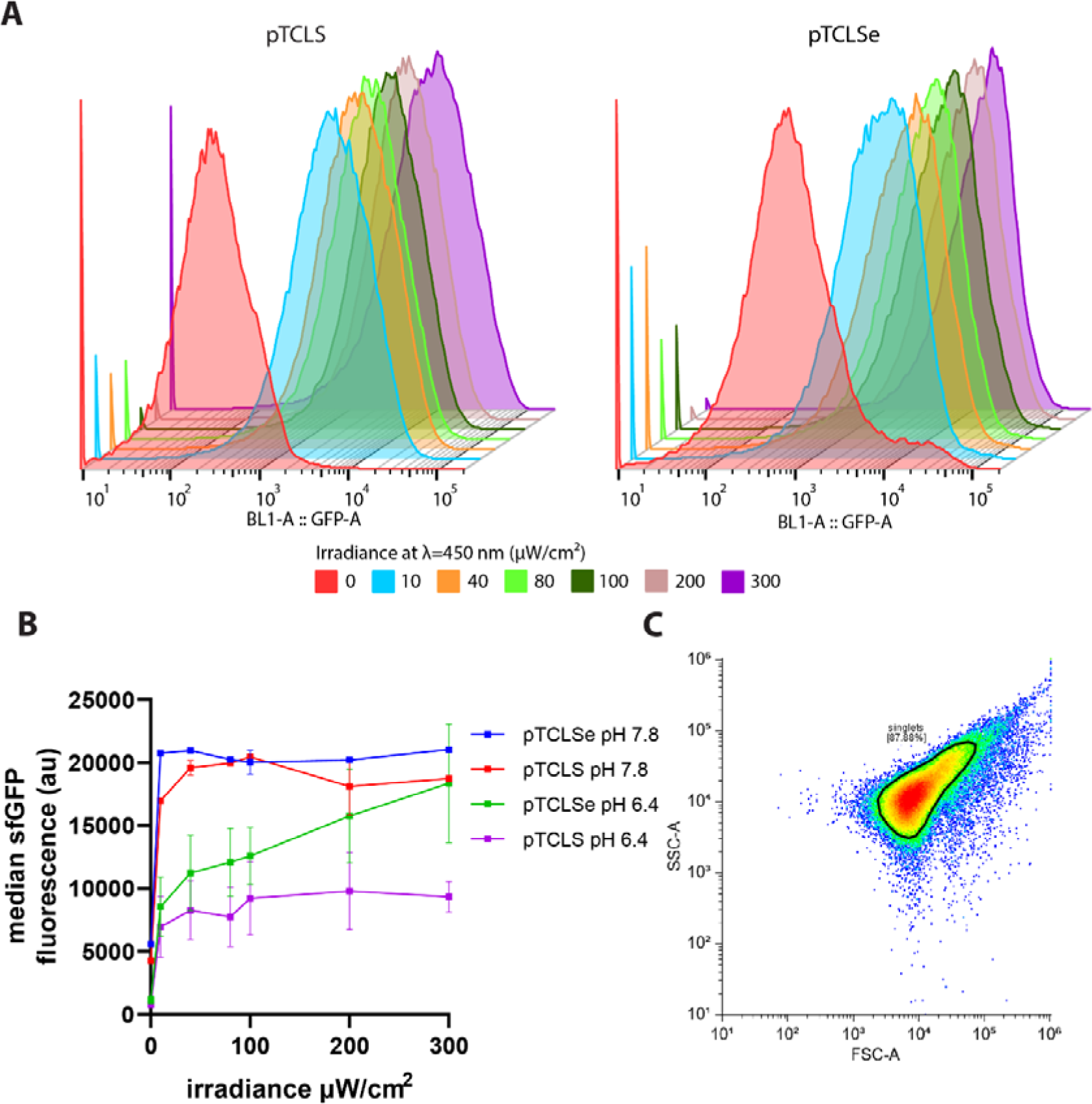
A) Representative flow cytometry histograms of *L. lactis* pTCLS (nMagHigh1/pMagFast2) and pTCLSe (eMagA/eMagB) irradiated with blue light for 5h at 0, 10, 40, 80, 100, 200 and 300 μW/cm² cultured in M17 supplemented with glucose/arginine. As the light intensity increases, the median increases monotonically, suggesting a strong correlation between the irradiance and the amount of expressed and detectable sfGFP. B) Scatter plot of the medians of the flow cytometry of three different experiments plotted against the incident light intensity. *L. lactis*-pTCLS seems to reach a saturating expression level of sfGFP after 80-100 μW/cm² while L. lactis-pTCLSe increases in a linear way up to 300 μW/cm² when cultured in M17 supplemented with glucose/arginine at 0.5% w/v, with an approximate final pH of 6.3. When cultured in M17 with 0.5% arginine (final pH ≈ 7.6), the T7RNAP is more active and the measured sfGFP intensity reaches a plateau even at lower irradiance values, 10 μW/cm² and an overall increased sfGFP expression level, with the drawback of a lower dynamic range since the basal expression levels are higher. Data shows the average ± SEM, n=3, of the median of independent FC experiments with at least 70,000 events recorded per experiment. C) FSC/SSC graph of a representative flow cytometry experiment showing the gating strategy, the aim was to select only individual cells and discard cells growing in chains, represented as cells that do not lie straight in the diagonal (comparable FSC and SSC values) of the scatter plot.

The eMags photoreceptors, while showing higher expression levels in the basal state at both low and high pH, can reach overall higher protein expression levels than the nMagHigh1/pMagFast2 variants, which require concatemerization and a 24 h pre-incubation step at 28°C to reach a mature folded state to become fully photoresponsive. Figure 3C illustrates the gating strategy used to discriminate individual *L. lactis* cells, since the gene expression patterns are highly heterogeneous within the population and measuring chains with cells with different sfGFP expression values would skew the data. Only events with comparable FSC/SSC values were analysed (figure 3C).

We next assessed the kinetics of sfGFP light-driven expression. *L. lactis* was cultured in GAM17C and AM17C (0.5% glucose/arginine and 0.5% arginine) starting from an overnight glycerinate stock grown in absence of light. Overnight precultures were diluted 1/5 in prewarmed GAM17C or AM17C medium and irradiated at 300 μW/cm² for up to 5h. An aliquot was taken every hour, cooled down immediately at 4°C, fixed in 4% formaldehyde as described before and analysed using flow cytometry. Results (figures 4A and 4B) reveal that the apparent sfGFP intensity starts at a low value at t=0h and gradually increases with time, reaching a maximum after 5h in all the tested conditions except for *L. lactis*-pTCLS cultured in glucose/arginine, where there is an apparent decrease in the fluorescence. It is worth noting that bacteria cultured in AM17C (pH ≈7.8) show a clear unimodal distribution in the flow cytometry histograms (figure 4A, lower row) while bacteria cultured in GAM17C start with a unimodal distribution of populations but in the samples taken at 4 and 5h, there is an apparent bimodal distribution of the population, with one of the peaks with an overall higher fluorescence than the samples taken earlier in the experiment and the other peak showing a diminished fluorescence than the average of the rest of the samples. Since the graph in figure 4B shows the average of the medians of the distribution for three independent experiments, this median is skewed towards a lower value, with an apparent overall lower intensity.

**Figure 4.**
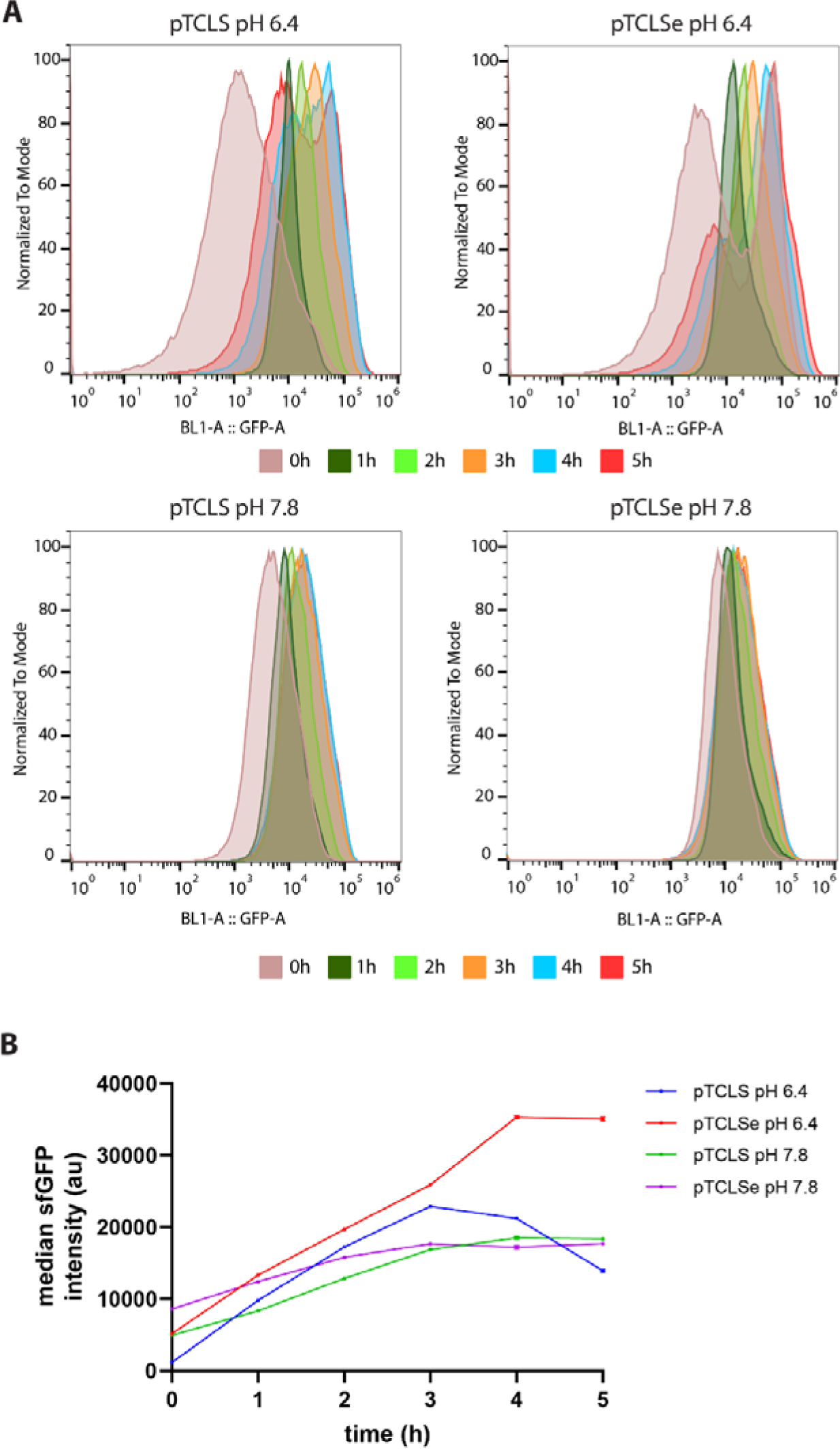
A) Representative flow cytometry histograms of *L. lactis*-pTCLS and pTCLSe cultured for up to 5h under blue light at 300 μW/cm² in M17 supplemented with either 0.5% glucose and 0.5% L-arginine (G/A) or only 0.5% arginine (A), with 10 μg/mL chloramphenicol in every condition. There is a clear correlation between the irradiation time and the overall sfGFP fluorescence as shown in B). It is interesting to note that *L. lactis* cultured in medium with glucose and arginine (pH 6.4) shows a bimodal distribution at 4 and 5h while the cells cultured in arginine (pH 7.8) reveal a unimodal distribution. Since the graph in B) describes the average and the standard deviation of the medians of three independent experiments, the apparent decline in intensity in the case of *L. lactis*-pTCLS cells cultured at pH 6.4 can be attributed to the displacement of the median to lower values due to the bimodal distribution. Overall, the fluorescence intensity increases with time suggesting a strong correlation between the irradiation time and the T7RNAP reconstitution and the subsequent expression of sfGFP. The standard deviation is too small to be shown in the graph.

We next wanted to determine if light-activated gene expression is active in *L. lactis* cells embedded in a hydrogel matrix, as one of the main aims of this work is the development a light-inducible gene expression system suitable for engineered living materials. We chose as a starting material Pluronic F-127, a hydrogel commonly used for bacterial encapsulation(*51–55*). Pluronic F-127 (PL from now on) is a poly(ethylene oxide)/poly(propylene oxide)/poly(ethylene oxide) (PEO–PPO–PEO) amphiphilic triblock copolymer with a lower critical solution temperature (LCST) that reversibly transitions from a low viscosity solution at low temperatures to a hydrogel at higher temperatures(*56*), becoming a fully crosslinked hydrogel at 30°C, the optimal growth temperature for *L. lactis*. In the gel phase, PL displays a shear-thinning behaviour that makes it appropriate for cold extrusion-based 3D printing, behaving as a viscous paste that keeps its structural integrity after extrusion. We firstly analysed the viability and proliferation of *L. lactis* embedded in GAM17C and AM17C mixed with 30% wt PL (Plu-GAM17C and Plu-AM17C from now on). Overnight *L. lactis* cultures were diluted 1/5 in cold Plu-GAM17C and Plu-AM17C and mixed to ensure a homogeneous suspension of the bacterial cells and cast in UV/vis cuvettes. The initial OD_590_ was in the range of 0.05-0.06 and the samples, three for pTCLS and pTCLSe in either Plu-GAM17C or Plu-AM17C were incubated at 30°C in airtight cuvettes for up to 8 days. Figure 5A shows the evolution of the optical density of the samples over time; in each tested condition, cells proliferate quickly during the first 24h and then the growth slows down and reaches a maximum after 5-6 days, suggesting that the physical hydrogel matrix slows down the proliferation of the bacterial cells even if there is still nutrient availability, since the matrix contains complete growth medium. It is worth noting that cells cultured in absence of glucose reach a lower final optical density than cells cultured with 0.5% glucose. This behaviour is promising since a common concern for the use of viable bacteria in engineered living materials is the proliferation control, which can be achieved using genetic elements (toxin-antitoxin systems, kill switches), bacteriostatics or physical entrapment in highly crosslinked matrices. We hypothesize here that embedding *L. lactis* in chemically crosslinked gels and suboptimal medium will hinder its proliferation rate even further.

**Figure 5.**
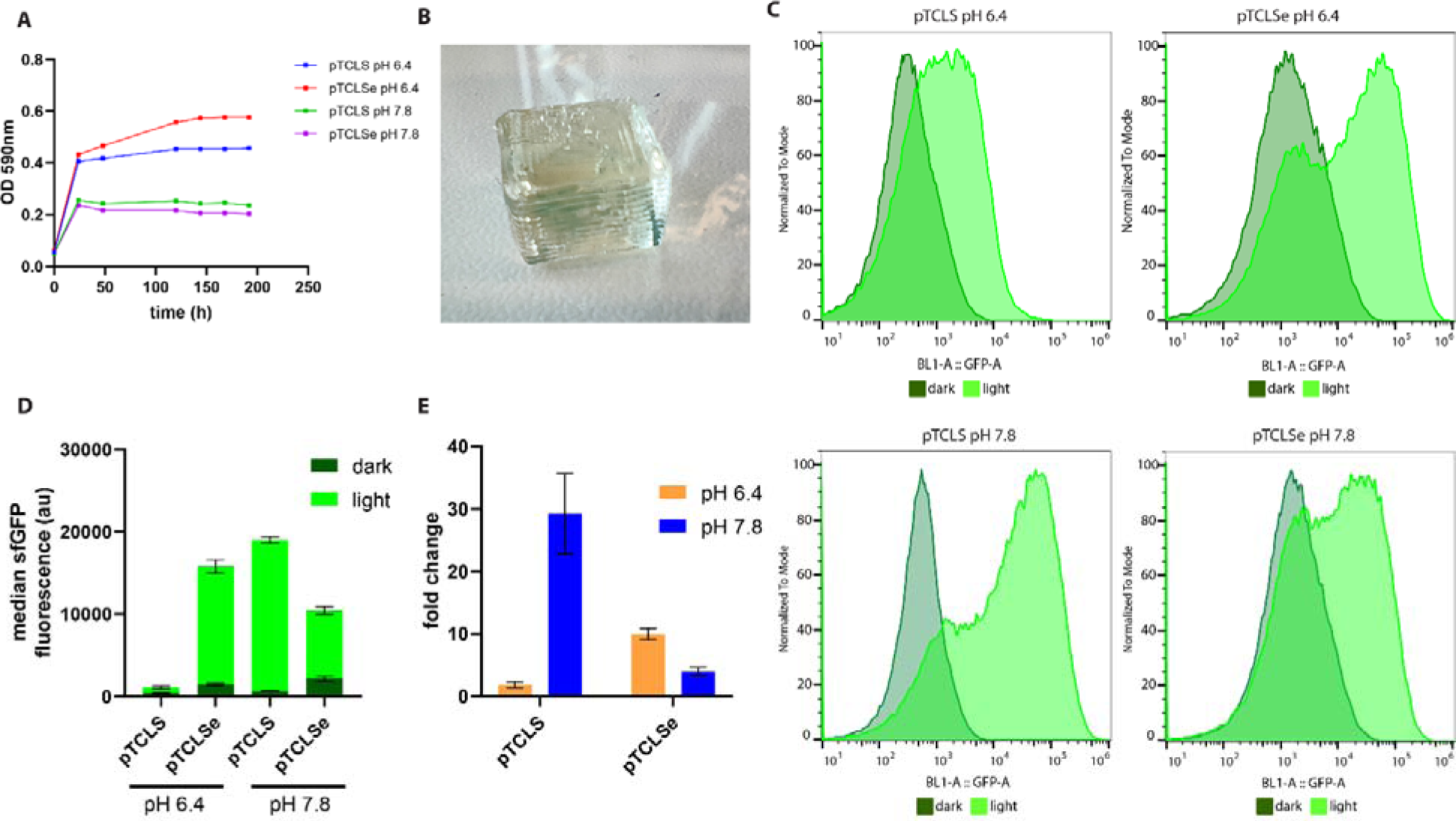
Light activation of *L. lactis* pTCLS and pTCLSe embedded in a 30% wt Pluronic F-127 hydrogel. A) *L. lactis*-pTCLS and pTCLSe cells with an initial OD_590_ density of 0.05-0.06 were embedded in Pluronic F-127 hydrogel with GAM17C or AM17C in microvolume UV/vis spectrophotometry cuvettes and incubated for up to 8 days, measuring the OD_590_ episodically. Apparently, there is an initial quick proliferation phase where cells reach a density of around 0.41-0.43 for the samples with glucose/arginine and 0.24-0.25 for samples with only 0.5% arginine. After 5-6 days, the proliferation stops, suggesting that the Pluronic matrix is able to contain the proliferation of the bacteria. B) Image of a 10×10×5 mm parallelepiped printed layer-by-layer in a Cellink BioX6 printer. The layers are clearly visible in the image. The OD_590_ of this sample is 0.05. C) Representative flow cytometry histograms (BL1 channel, corresponding to sfGFP) of bacterial samples. There is a clear activation driven by the blue light in each condition, strongly suggesting that blue light can deeply penetrate in the optically transparent hydrogel and drive the T7RNAP dimerization, transcription and sfGFP expression even in slowly proliferating cells physically confined in a hydrogel. D) Activation values (average the median of 3 independent experiments) corresponding to cells cultured in absence or with blue light with an irradiance of 300 μW/cm² for 5h. The activation values depend on the strain and the condition. E) Fold change values derived from graph D, again the fold change values depend on the condition, being *L. lactis*-pTCLS grown in Plu-GAM17C the condition with the lowest fold change and activation. It is interesting to note that *L. lactis*-pTCLS grow in Plu-AM17C showed a notable fold change and overall activation levels.

We wanted next to assess if *L. lactis* physically embedded in PL-M17 hydrogels would still respond to blue light and express sfGFP. As mentioned above, a 30% wt Pluronic hydrogel mixed with GAM17C or AM17C at an initial OD_590_ of 0.05-0.06 (approximately 5 to 6×10□ CFU/mL) was used to 3D print 10×10×5 mm parallelepipeds using a Cellink BioX6 bioprinter (Figure 5B). The constructs were printed under red light to avoid unwanted activation by blue light for the “dark” controls. The samples were then transferred to an airtight container and irradiated with blue light at 300 μW/cm² for 5h, in a 30°C incubator oven. After that time, cells were recovered from the hydrogel (see the relevant materials and methods section) and analysed using flow cytometry. As shown in figure 5C and 5D, we observed significant light activation in terms of sfGFP expression in all the tested conditions. Even in a lower proliferating state, cells were still able to sense and respond to the blue light but in this case the results were slightly different to cells cultured in the log phase in liquid medium. *L. lactis*-pTCLS grown in GAM17C showed the lower fold change in activation (figure 5E), around 2, while pTCLS in AM17C and pTCLSe in GAM17C showed the highest fold change values. Even if the differences between conditions are appreciable, there is still an unambiguous level of activation of the T7RNAP-driven transcription/expression machinery. This result warrants a deeper look in this direction, embedding the cells in chemically crosslinked hydrogels and suboptimal growth medium.

## Discussion

In the present study we show the development of an optogenetic gene expression system based on a split T7 RNA polymerase fused to the blue light-sensitive engineered heterodimerizing Magnets and an enhanced version, derived from *Neurospora crassa.* The T7 RNA polymerase is a highly important enzyme widely used in synthetic biology and biotechnology for its desirable characteristics, being a single subunit enzyme that does not depend on sigma factors and can be split and reconstituted while maintaining its activity.

We show how an optogenetic system based on a split T7 RNA polymerase and heterodimerizing photoreceptors can be successfully adapted to *Lactococcus lactis*, an important gram-positive lactic acid bacterium. This is, to the best of our knowledge, the second optogenetic system developed for *L. lactis*(57) with the advantage that the present system is orthogonal to the native bacterial expression machinery and can be decoupled using for example rifampicin, an inhibitor of the native RNA polymerase that does not affect the T7RNAP. The present system can be further optimized by adjusting the expression levels of the N-terminal and C-terminal fragments of the T7RNAP, by using promoters of variable potency or engineered RBS. We use constitutive promoters that are always active thus avoiding the dependency on chemically inducible promoters, but we are aware that for some applications it might be more appropriate to express the T7RNAP fragments in an inducible way, during only certain phases of the cell culture, to further restrict the low-level basal protein expression observed throughout this work. Moreover, the ratio between the two fragments of the T7RNAP can be adjusted as previously described(*21*) to tune the basal and activated gene expression in response to blue light.

During this study, a new work was published describing the development of an enhanced version of the Magnet photoreceptors. The variant used in the pTCLS plasmid requires a 24h pre-incubation step at 28°C to allow for the complete folding and maturation of the photoreceptors and the dimerization efficiency is not optimal, requiring the concatemerization of the receptors to achieve an optimal response to blue light. The receptors are also not suitable for its use at 37°C due to their thermolability, which difficult its use in mammalian cells. The new version or “enhanced Magnets” overcome those limitations by allowing its use at 37°C and achieving a higher heterodimerization efficiency using single subunits of the photoreceptors.

We developed two variants of the optogenetic system using both the older and the newer versions, showing that both are active in *L. lactis*. According to our data, the system with the enhanced Magnets shows an overall higher gene expression efficiency when the cells are in the log phase of the growth in liquid medium at the cost of a reduced dynamic range, while the classic Magnets show an increased dynamic range with an overall lower expression efficiency. This system is also active when the cells are embedded in a physically crosslinked hydrogel, Pluronic F-127 in our case. *L. lactis* still responds to blue light by expressing sfGFP even if the proliferation rate is greatly slowed down, which is expected due to the orthogonality of the T7RNAP respect to the native bacterial machinery.

The well-known pH-dependent activity of the T7RNAP is also evident in this work. Lactic acid is the main metabolite of the homofermentative glycolytic pathway used by *L. lactis* to synthesize ATP; being a weak organic acid, the environmental pH quickly lowers to values around 5.5, which means that in the later stages of a *L. lactis* cell culture with glucose present in the medium, the T7RNAP activity will be greatly reduced or even suppressed. We show an easy strategy to tackle this issue by taking advance of the arginine deiminase pathway, which metabolizes L-arginine to CO_2_, NH_3_ and L-ornithine. L-arginine supplementation in the medium allows the production of ammonia, that counteracts the acidity of lactic acid shifting the pH to values between 7 and 8, restoring the activity of the T7RNAP while maintaining the viability of *L. lactis*.

The two systems presented here is blue-light responsive, but since the Magnets are only a part of the whole light-controlled expression machinery, they could be potentially exchanged with other photoreceptors that respond to different wavelengths. Recently developed photoreceptors include BICYCLs(*58*), sensitive to green or red light, and MagRed(*59*), sensitive to red light, could be used to develop a full multichromatic expression system.

The potential applications for the present system go beyond living materials; *L. lactis*, with its effective secretion systems, can be an adequate substitute for *E. coli* or *V. natriegens* while lacking lipopolysaccharide expression, enabling the production of higher quality recombinant proteins. The use of a light-induced expression system allows to decouple the growth and production phases using only light, an economically viable alternative to more expensive inducers such as nisin.

## Materials and methods

### *L. lactis* cell culture

*L. lactis* cells were cultured using M17 medium (*60*) supplemented with 0.5% wt glucose and 0.5% wt L-arginine (GAM17C), 0.5% L-arginine (AM17C) with 10 μg/mL chloramphenicol in anaerobic conditions as standing cultures at 30°C without shaking, although it tolerates mild aerobic and shaking conditions.

Agar plates for transformation were prepared mixing 1.5% agar in M17 supplemented with 0.5% glucose and the relevant antibiotic. The plates were incubated at 30°C overnight.

### Molecular cloning and plasmid construction

The genetic sequences of the T7RNAP R632S, nMagHigh1, pMagFast2, eMagA, eMagB, CPxx synthetic promoters and RBS were either amplified from plasmids acquired from Addgene, from our stocks, or purchased as synthetic DNA fragments from Twist Bioscience or Genscript.

Constructs were designed *in silico* in Benchling and primers for the Gibson reaction were obtained from the NEBuilder website (New England Biolabs). Primers were purchased from Sigma-Aldrich in cartridge purified quality and used in PCR reactions with the Q5 high fidelity (NEB) or the Platinum SuperFi II polymerase (ThermoFisher). PCR products were purified using a Qiagen PCR purification kit and assembled using the NEBuilder HiFi DNA assembly kit following the manufacturer’s instructions.

The constructs were then transformed in *L. lactis* as described in the bacterial transformation section. The recombinant constructs were analysed by agarose gel electrophoresis to confirm their size and sequenced using Oxford nanopore technology (Full Circle Labs Bio, UK).

### Bacterial transformation

Electrocompetent *L. lactis* NZ9000 cells were prepared according to Holo and Nes(*61*). In summary, *L. lactis* cells are precultured overnight in GM17 supplemented with 1.5% wt glycine. The preculture is then diluted 1/100 in fresh GM17 supplemented with 1.5% glycine and 0.5 M sucrose and grown at 30°C until OD_590_ 0.2-0.3. Then, the culture is quickly cooled down to 4°C and washed three times with washing buffer (ice-cold 0.5M sucrose and 10% glycerol in sterile ultrapure water). The first wash is done in the original volume, the second in ½ volumes, the third in ⅓ volume and the final pellet is resuspended in 1% of the original volume culture. All the centrifugations are performed on a tabletop centrifuge at 6000-7000g and 4°C. The resulting suspension is aliquoted in volumes of 50 μL, snap-frozen in liquid nitrogen and stored at -80°C or used immediately.

The electrotransformation is performed by thawing electrocompetent cells for 10 min in ice, mixing 1 μL of the purified plasmid or assembly reaction with 50 μL of the electrocompetent cells, which are transferred to a 1 mm gap electroporation cuvette previously chilled to 4°C and immediately electroporated using a 2 kV pulse, with a typical time constant of 4.8-5 ms. Cells are immediately resuspended in 950 μL of recovery medium (ice-cold M17 supplemented with 0.5% glucose, 2 mM CaCl□ and 20 mM MgCl□) and incubated for 2h at 30°C. The culture is centrifuged at 7000 g for 3 min, the pellet resuspended in 100 μL recovery medium and plated in agar 1.5%-GM17 plates with the appropriate antibiotic for 24h at 30°C.

### Blue light-induced gene expression

All the experiments were performed in triplicate unless otherwise stated. Overnight *L. lactis* precultures from glycerol stocks originated from single colonies were grown in vessels protected from light at 30°C in G/AM17C medium. These cultures were then diluted in variable proportions in fresh G/AM17C prewarmed to 30°C. Uninduced bacteria were grown in P24 multiwell plates wrapped in aluminium foil while induced bacteria were cultured a homemade 3D-printed opaque PLA P24 multiwell plate with a transparent polymeric polyolefine plastic film bottom located on top of a homemade device that illuminates each individual well in the plate from the bottom using an in-house fabricated device. This device consists of a Feather M4 development board (Adafruit Inc.) programmed with circuit python and connected to a 24-channel TLC5947 PWM LED controller with 12-bit output (4096 values) wired to an array of 24 individual blue LEDs (emission maximum at 470 nm). A python script was used to control the intensity and timing of each individual addressable LED. The LED intensities were measured and calibrated to an average coefficient of variation of the output intensity of 1.8% or better, allowing for adjustable outputs from 0 to 2000 μW/cm² in 0.48 μW/cm² increments. The board and software allow for a timing controlled down to the millisecond.

All experiments were performed inside a dry incubator oven protected from light. *L. lactis* cells were collected at variable time points, depending on the experiment, and immediately centrifuged at 12 000g and 4°C for 2 min to stop the cell metabolism and the activity of the T7 RNA polymerase. The pellet was then resuspended in cold 4% formaldehyde in PBS and fixed for 15 min at room temperature. Cells were then centrifuged at 12 000g for 4 minutes and the cell pellet was resuspended in PBS and kept in the dark at 4°C until further analysis, either fluorescence readings in the plate reader or flow cytometry measurements. Fluorometry and flow cytometry analyses were performed within 24h of the light activation experiment.

### Flow cytometry

After the culture and light irradiation, *L. lactis* cells were washed by centrifugation at 12 000g for 1 minute and resuspended in an equal volume of PBS, left overnight at 4C to ensure that the sfGFP was fully mature, and then subjected to a 2-minute, 50 Hz cycle in a Qiagen TissueLyser homogeneizer to break down bacterial chains. The cell suspension was analyzed in a ThermoFisher Attune NxT flow cytometer with a FSC voltage of 380V, SSC voltage of 420 V and a BL1 (488 nm blue laser) gain of 250V. The cytometer was instructed to acquire a total of 100 000 events at 25 μL/min. The resulting data was exported as FCS files and analyzed with FlowJo VX 10.

The FSC/SSC scatter plots were gated to ensure that only individual cells were analyzed, and then the data was plot as histograms with the BL1-area values for sfGFP fluorescence and plotted using a logarithmic graph with the y axis normalized to mode. FlowJo VX 10 was used to calculate and display the statistics for the histograms.

### 3D bioprinting and encapsulation

*L. lactis* cells were encapsulated in a 30% wt solution of Pluronic F127 in either GAM17C or AM17C. Firstly, M17 media was prepared and autoclaved, then supplemented with glucose/arginine or arginine and chloramphenicol and cooled down, followed by the addition of Pluronic F127. The solution was kept at 4°C under agitation overnight. The following day, overnight cultures of *L. lactis*-pTCLS and pTCLSe grown in AM17C in absence of light were concentrated by a factor of ten and a volume equivalent to 1/5 of the original undiluted culture was mixed with cold Pluronic F-127 30% solution and kept at 4°C in a tube roller for 20 minutes. The initial OD_590_ values for the Pluronic solution ranged between 0.05 and 0.07. The cold Pluronic/*L. lactis* mixes were transferred to 3 mL printer cartridges and kept for 10 minutes at 30°C in an oven incubator to allow for the physical crosslinking of the hydrogel.

After that time, the constructs were transferred to a centrifuge tube and cooled down to 4°C to stop the bacterial metabolism and allow the hydrogel to go back to the liquid sol phase; after 24h, the cells were centrifuged at 10,000 g for 10 minutes to pellet the bacteria, the viscous solution was removed with a pipette and the cells were resuspended again in cold PBS to ensure the total resuspension of the cells.

Cells were then collected by centrifugation, fixed and analysed using flow cytometry as described in the relevant section.

## Acknowledgements

Plasmid pN565 was a gift from Christopher Voigt (Addgene plasmid # 49990; http://n2t.net/addgene:49990; RRID:Addgene_49990)

