## Supplementary figures and table for "Development of an optogenetic gene expression system in *Lactococcus lactis* using a split photoactivatable T7 RNA polymerase"

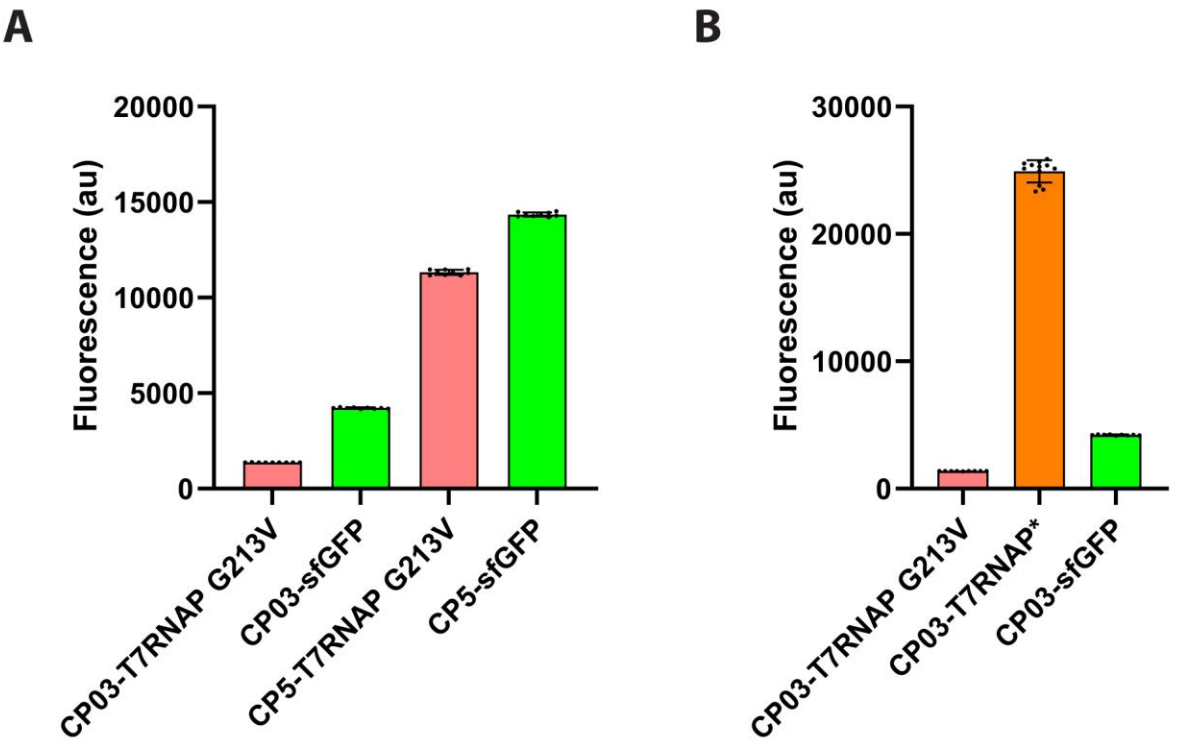


**Supplementary figure 1**. A) Effect of the G213V mutation on the T7RNAP. Fluorometry measurements of *L. lactis* transformed with a plasmid containing the T7 RNA polymerase under the control of either CP03 or CP5 promoter. The expressed T7RNAP drives the expression of sfGFP located downstream under the control of an L. lactis-optimized T7 promoter and RBS (green bars) compared to the direct expression of sfGFP under the same promoters. The effect of the G213V mutation is clear in the sense of keeping its transcriptional activity but at lower levels. B) Comparison of the G213V mutated T7RNAP and the variant used in the rest of this work, which only contains the R632S mutation, and its comparison with the expression of sfGFP under the direct control of the CP03 promoter.

A

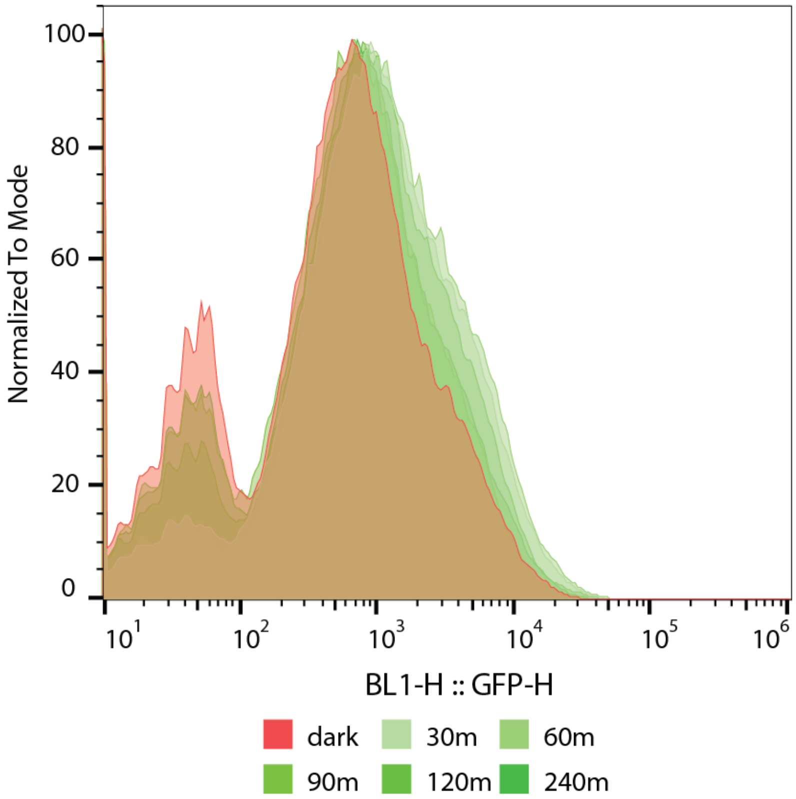


**Supplementary figure 2**. sfGFP photobleaching. To assess whether the use of blue light, which is needed to heterodimerize the Magnets, would have an effect on the fluorescence of the sfGFP, we measured the fluorescence of *L. lactis* expressing sfGFP under the control of the CP5 promoter. An overnight L. lactis culture was split in two and one aliquot was kept protected from light (red histogram) for 4h while another aliquot was irradiated with a 470 nm LED at 100 $\mu W/cm^2$ for 30, 60, 90, 120 and 240 minutes respectively. Cells were homogenised in a tissue disruptor shaker, centrifuged, resuspended in PBS and analyzed in an Attune NxT flow cytometer. We can infer from the histograms that there is no appreciable photobleaching after 4h under constant medium intensity blue light, since the control bacteria shows a very similar intensity histogram without any noticeable reduction in intensity. In fact, and probably due to the inherent variability of the flow cytometry, the sample kept protected from light has a slightly lower overall intensity in terms of median and robust CV as shown in the table. Data suggests that any difference observed during the light activation experiments corresponds only to the different sfGFP expression levels and that photobleaching during the experiment is negligible.

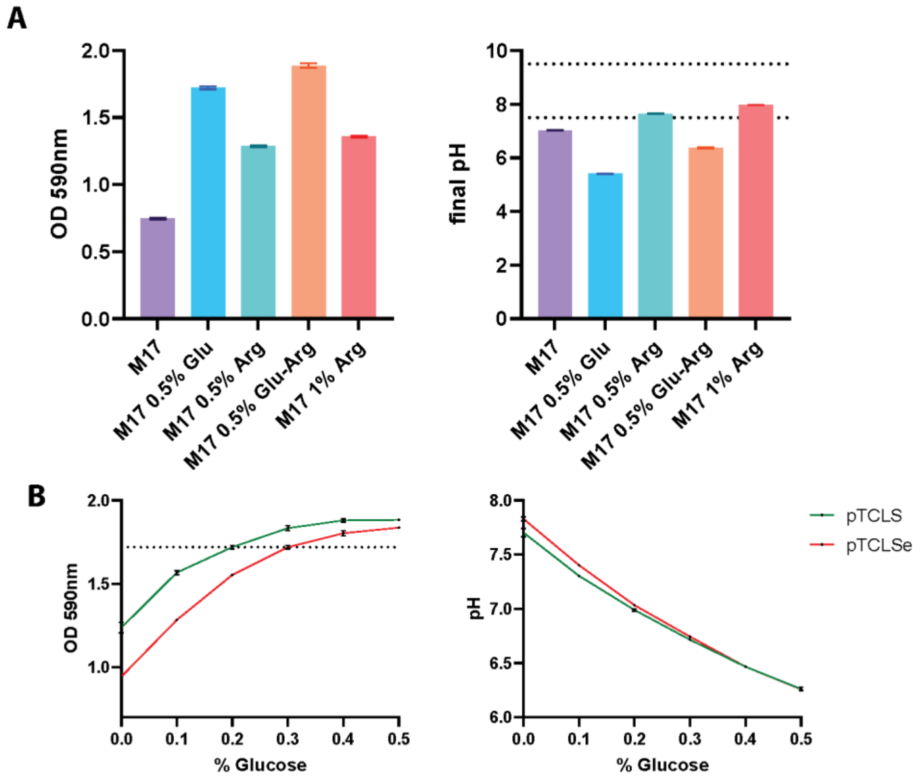


**Supplementary figure 3**. Effect of the media composition on the final $OD_{590}$ and the pH of the medium. The activity of the T7 RNA polymerase is highly dependent on the pH(*1*) with an optimum between pH 7.5 and 9.5. *L. lactis* main metabolite in presence of glucose is lactic acid, that lowers the pH of the medium. An easy strategy to reverse this effect is to take advantage of the arginine deiminase pathway, that uses L-arginine to obtain ATP producing ornithine, $CO_2$ and $NH_3$ as the main catabolites. A) Optical density and pH of *L. lactis* cultures grown in M17 without supplements and M17 supplemented with 0.5% glucose, 0.5% arginine, 0.5% glucose and 0.5% arginine or 1% arginine. The biomass produced in absence of glucose is lower compared to cells cultured in presence of glucose, but the presence of arginine still increases the overall $OD_{590}$ compared to bare M17 medium. The final pH of the cultures with arginine and without glucose reached the minimum value of 7.5 needed for a sufficiently active T7RNAP. B) Effect of smaller amounts of glucose on biomass production and pH on the two optogenetic systems discussed here, pTCLS and pTCLSe. Cells were cultured in M17 supplemented with 0.5% arginine and variable concentrations of glucose from 0 to 0.5%. Small concentrations of glucose had a noticeable effect on the final pH and biomass production; even 0.1% glucose led to a pH diminution from approximately 7.7-7.8 to 7.3-7.4 and went down to pH 7 at 0.2% glucose, with the corresponding decrease on the activity of the T7RNAP. The increment in the biomass from 0 to 0.2% glucose was notable as well, but ODs of around 0.9 to 1.2 are more than enough for our application and what is most important, the optogenetic system is active.

**Supplementary table 1**. Genetic elements used throughout this work.

| | |
|---|---|
| Promoter CP03 | CATGGGTAAGTTTATTCTTCACACTATCTGGGCCCGATGGTATAATAAGTGACT<br>ACTGTT |
| Promoter CP2 | CATCATTAAGTTTATTCTTCACATTGGCCGGAATTGTTGTATAATACCTTAGTA<br>CTGTT |
| Promoter CP5 | GATGTTTTAGTTTATTCTTGACACCGTATCGTGCGCGTGATATAATCGGGATCC<br>TTAAGA |
| Promoter CP12 | CATCGGGTAGTTTATTCTTGACAATTAAGTAGAGCCTGATATAATAGTTCAGTA<br>CTGTT |
| Promoter CP50 | CATCGCTTAGTTTTTCTTGACAGGAGGGATCCGGGTTGATATAATAGTTAGTAC<br>TGTT |
| Promoter CP1000 | TATGCGGTAGTTTATTCTTGACATGACGAGACAGGTGTGGTATAATGGGTCTAG<br>ATTAGG |
| 5’ UTR of T7g10 gene and RBS | GGGAGACCACAACGGTTTCCCACTAGAAATAATTTTGTTTAACTTTAGAAAGGA<br>GATATACGC |
| N-terminal end of T7RNAP<br><br>Start codon | ATGAACACGATTAACATCGCTAAGAACGACTTCTCTGACATCGAACTGGCTGCT<br>ATCCCGTTCAACACTCTGGCTGACCATTACGGTGAGCGTTTAGCTCGCGAACAG<br>TTGGCCCTTGAGCATGAGTCTTACGAGATGGGTGAAGCACGCTTCCGCAAGATG<br>TTTGAGCGTCAACTTAAAGCTGGTGAGGTTGCGGATAACGCTGCCGCCAAGCCT<br>CTCATCACTACCCTACTCCCTAAGATGATTGCACGCATCAACGACTGGTTTGAG<br>GAAGTGAAAGCTAAGCGCGGCAAGCGCCCGACAGCCTTCCAGTTCCTGCAAGAA<br>ATCAAGCCGGAAGCCGTAGCGTACATCACCATTAAGACCACTCTGGCTTGCCTA<br>ACCAGTGCTGACAATACAACCGTTCAGGCTGTAGCAAGCGCAATCGGTCGGGCC<br>ATTGAGGACGAGGCTCGCTTCGGTCGTATCCGTGACCTTGAAGCTAAGCACTTC<br>AAGAAAAACGTTGAGGAACAACTCAACAAGCGCGTAGGGCACGTCTACAAGAAA<br>GCATTTATGCAAGTTGTCGAGGCTGACATGCTCTCTAAGGGTCTACTCGGTGGC<br>GAGGCGTGGTCTTCGTGGCATAAGGAAGACTCTATTCATGTAGGAGTACGCTGC<br>ATCGAGATGCTCATTGAGTCAACCGGAATGGTTAGCTTACACCGCCAAAATGCT<br>GGCGTAGTAGGTCAAGACTCTGAGACTATCGAACTCGCACCTGAATACGCTGAG<br>GCTATCGCAACCCGTGCAGGTGCGCTGGCTGGCATCTCTCCGATGTTCCAACCT<br>TGCGTAGTTCCTCCTAAGCCGTGGACTGGCATTACTGGTGGTGGCTATTGGGCT<br>AACGGTCGTCGTCCTCTGGCGCTGGTGCGTACTCACAGTAAGAAAGCACTGATG<br>CGCTACGAAGACGTTTACATGCCTGAGGTGTACAAAGCGATTAACATTGCGCAA<br>AACACCGCATGGAAAATCAACAAGAAAGTCCTAGCGGTCGCCAACGTAATCACC<br>AAGTGGAAGCATTGTCCGGTCGAGGACATCCCTGCGATTGAGCGTGAAGAACTC<br>CCGATGAAACCGGAAGACATCGACATGAATCCTGAGGCTCTCACCGCGTGGAAA<br>CGTGCTGCCGCTGCTGTGTACCGCAAGGACAAGGCTCGCAAGTCTCGCCGTATC<br>AGCCTTGAGTTCATGCTTGAGCAAGCCAATAAGTTTGCTAACCATAAGGCCATC<br>TGGTTCCCTTACAACATGGACTGGCGCGGTCGTGTTTACGCTGTGTCAATGTTC<br>AACCCGCAAGGTAACGATATGACCAAAGGACTGCTTACGCTGGCGAAAGGTAAA<br>CCAATCGGTAAGGAAGGTTACTACTGGCTGAAAATCCACGGTGCAAACTGTGCG<br>GGTGTCGACAAGGTTCCGTTCCCTGAGCGCATCAAGTTCATTGAGGAAAACCAC<br>GAGAACATCATGGCTTGCGCTAAGTCTCCACTGGAGAACACTTGGTGGGCTGAG<br>CAAGATTCTCCGTTCTGCTTCCTTGCGTTCTGCTTTGAGTACGCTGGGGTACAG<br>CACCACGGCCTGAGCTATAACTGCTCCCTTCCGCTGGCGTTTGACGGGTCTTGC<br>TCTGGCATCCAGCACTTCTCCGCGATGCTCCGAGATGAGGTAGGTGGTCGCGCG<br>GTTAACTTGCTTCCT |
| C-terminal end of T7RNAP<br><br>Stop codon | AGTGAAACCGTTCAGGACATCTACGGGATTGTTGCTAAGAAAGTCAACGAGATT<br>CTACAAGCAGACGCAATCAATGGGACCGATAACGAAGTAGTTACCGTGACCGAT<br>GAGAACACTGGTGAAATCTCTGAGAAAGTCAAGCTGGGCACTAAGGCACTGGCT<br>GGTCAATGGCTGGCTTACGGTGTTACTCGCAGTGTGACTAAGAGTTCAGTCATG<br>ACGCTGGCTTACGGGTCCAAAGAGTTCGGCTTCCGTCAACAAGTGCTGGAAGAT<br>ACCATTCAGCCAGCTATTGATTCCGGCAAGGGTCTGATGTTCACTCAGCCGAAT<br>CAGGCTGCTGGATACATGGCTAAGCTGATTTGGGAATCTGTGAGCGTGACGGTG<br>GTAGCTGCGGTTGAAGCAATGAACTGGCTTAAGTCTGCTGCTAAGCTGCTGGCT |

| | |
|---|---|
| | GCTGAGGTCAAAGATAAGAAGACTGGAGAGATTCTTCGCAAGCGTTGCGCTGTG<br>CATTGGGTAACTCCTGATGGTTTCCCTGTGTGGCAGGAATACAAGAAGCCTATT<br>CAGACGCGCTTGAACCTGATGTTCCTCGGTCAGTTCCGCTTACAGCCTACCATT<br>AACACCAACAAAGATAGCGAGATTGATGCACACAAACAGGAGTCTGGTATCGCT<br>CCTAACTTTGTACACAGCCAAGACGGTAGCCACCTTCGTAAGACTGTAGTGTGG<br>GCACACGAGAAGTACGGAATCGAATCTTTTGCACTGATTCACGACTCCTTCGGT<br>ACGATTCCGGCTGACGCTGCGAACCTGTTCAAAGCAGTGCGCGAAACTATGGTT<br>GACACATATGAGTCTTGTGATGTACTGGCTGATTTCTACGACCAGTTCGCTGAC<br>CAGTTGCACGAGTCTCAATTGGACAAAATGCCAGCACTTCCGGCTAAAGGTAAC<br>TTGAACCTCCGTGACATCTTAGAGTCGGACTTCGCGTTCGCGTAA |
| T7 promoter adapted to *L. lactis*<br><br>T7 Promoter<br>Start codon | CCCAATACGCAAACCGCCTCTCCCCGCGCGTTGGCCGATTCATTAATGCAGGAT<br>CTCGATCCCGCGAAATTAATACGACTCACTATAGGGAGACCACAACGGTTTCCC<br>TCTAGAAATAATTTTGTTTAACTTTAAGAAGGAGATATACATATGCGGGGTTCT<br>CATCATCATCATCATCATGGTATGGCTAGCATGACTGGTGGACAGCAAATGGGT<br>CGGGATCTGTACGACGATGACGATAAG |
| nMagHigh1<br><br>Stop codon | CACACTCTTTACGCCCCTGGAGGATACGACATTATGGGATATTTGGATCAGATT<br>GGGAACCGCCCAAACCCTCAGGTCGAACTGGGGCCTGTGGACACGTCATGTGCC<br>CTGATCCTGTGCGATCTGAAGCAAAAGGACACTCCGATCGTCTACGCCTCGGAA<br>GCCTTCTTGTATATGACCGGATACAGCAATGCAGAGGTGCTCGGCAGGAACTGC<br>AGATTCCTGCAGTCCCCCGACGGGATGGTGAAACCAAAGTCGACTCGCAAATAT<br>GTGGACTCGAACACGATCAACACCATCCGGAAGGCCATCGACCGGAACGCCGAG<br>GTCCAGGTGGAGGTGGTCAACTTTAAGAAGAACGGCCAGCGGTTCGTGAACTTT<br>CTGACCATCATTCCGGTCCGGGATGAAACCGGAGAGTACAGATACTCCATGGGA<br>TTCCAGTGCGAAACCGAATAA |
| pMagFast2<br><br>Start codon | ATGCACACTCTTTACGCCCCTGGAGGATACGACATTATGGGATATTTGCGGCAG<br>ATTAGGAACCGCCCAAACCCTCAGGTCGAACTGGGGCCTGTGGACACGTCATGT<br>GCCCTGGTCCTGTGCGATCTGAAGCAAAAGGACACTCCGGTGGTCTACGCCTCG<br>GAAGCCTTCTTGTATATGACCGGATACAGCAATGCAGAGGTGCTCGGCAGGAAC<br>TGCAGATTCCTGCAGTCCCCCGACGGGATGGTGAAACCAAAGTCGACTCGCAAA<br>TATGTGGACTCGAACACGATCAACACCATGCGGAAGGCCATCGACCGGAACGCC<br>GAGGTCCAGGTGGAGGTGGTCAACTTTAAGAAGAACGGCCAGCGGTTCGTGAAC<br>TTTCTGACCATGATTCCGGTCCGGGATGAAACCGGAGAGTACAGATACTCCATG<br>GGATTCCAGTGCGAAACCGAA |
| eMagA<br><br>Start codon | ATGGGTCACACACTATACGCACCAGGTGGTTATGATATTATGGGTTATCTTGAT<br>CAAATCGCTAACAGACCAAACCCACAAGTTGAATTAGGACCAGTTGACCTGTCT<br>TGTGCTCTTATCTTATGTGACCTTAAACAAAAAGATACACCAATCGTTTATGCC<br>TCAGAAGCTTTTTTAGAAATGACTGGTTACAATCGTCATGAAGTACTCGGTCGT<br>AACTGTCGCTTTTTACAATCACCTGACGGTATGGTTAAACCTAAAAGCACCCGT<br>AAATATGTAGACTCTAACACAATCTATACAATTAAAAAAGCAATTGATCGTAAC<br>GCTGAAGTACAAGTTGAAGTTGTTAACTTTAAAAAAAATGGTCAACGTTTCGTC<br>AATTTTCTTACAATTATCCCAGTACGCGACGAAACTGGTGAATACCGATACTCA<br>ATTGGATTCCAATGTGAAACAGAA |
| eMagB<br><br>Stop codon | ATGGGACACACGCTTTACGCACCTGGTGGTTATGATATTATGGGTTATTTACGT<br>CAAATTCGCAACCGTCCAAACCCTCAAGTTGAACTTGGACCTGTAGACTTATCT<br>TGTGCACTCGTTTTATGTGACTTGAAACAAAAAGACACACCAGTTGTTTACGCA<br>TCAGAAGCTTTTTTAGAAATGACTGGATATAACCGTCATGAAGTACTTGGTCGT<br>AATTGCCGTTTTTTACAATCTCCAGATGGTATGGTTAAACCAAAGTCAACACGT<br>AAATACGTGGATTCAAACACAATTTACACTATGAAAAAAGCTATTGATCGTAAT<br>GCTGAAGTTCAAGTCGAAGTTGTAAATTTTAAAAAAAATGGTCAACGTTTCGTT<br>AATTTCCTCACAATGATCCCAGTACGAGATGAAACTGGTGAATACCGTTACTCT<br>ATCGGATTCCAATGTGAAACAGAATAA |
| GGSGG Linker | GGCGGTTCTGGAGGT |
| Superfolder GFP<br><br>Start codon | ATGTCAAAAGGAGAAGAGCTGTTCACAGGTGTTGTGCCGATTCTCGTTGAGCTT<br>GACGGAGATGTAAACGGACACAAATTCTCTGTTCGCGGTGAAGGTGAAGGAGAT<br>GCAACAAACGGCAAGCTGACATTGAAGTTTATTTGCACAACTGGAAAGCTGCCG<br>GTTCCTTGGCCGACACTTGTAACGACGCTGACTTACGGCGTTCAATGCTTCTCT |

| | |
|---|---|
| Stop codon | CGTTATCCAGACCACATGAAACGCCATGATTTCTTCAAATCTGCAATGCCTGAA<br>GGCTACGTTCAAGAGCGTACGATCAGCTTCAAAGATGACGGAACGTACAAAACA<br>AGAGCAGAAGTGAAGTTTGAAGGTGACACACTTGTGAACCGCATTGAATTGAAA<br>GGCATTGATTTCAAAGAAGATGGAAACATCCTTGGACACAAACTTGAATACAAC<br>TTCAACAGCCACAACGTATACATCACTGCTGACAAACAAAAAAACGGCATCAAA<br>GCAAACTTCAAAATCCGTCATAACGTAGAGGACGGTTCTGTTCAGCTTGCTGAT<br>CATTATCAGCAAAATACACCGATCGGTGACGGCCCGGTTCTTCTTCCTGATAAC<br>CATTATTTATCAACTCAAAGCGTATTATCAAAAGACCCAAATGAAAAGCGTGAC<br>CACATGGTGCTGCTTGAATTTGTGACAGCTGCTGGTATCACTCACGGCATGGAT<br>GAGCTTTATAAGTAA |
